# Human embryonic stem cells-derived dopaminergic neurons transplanted in parkinsonian monkeys recover dopamine levels and motor behavior

**DOI:** 10.1101/2020.07.08.192591

**Authors:** Adolfo López-Ornelas, Itzel Escobedo-Avila, Gabriel Ramírez-García, Rolando Lara-Rodarte, César Meléndez-Ramírez, Beetsi Urrieta-Chávez, Tonatiuh Barrios-García, Verónica A. Cáceres-Chávez, Xóchitl Flores-Ponce, Francia Carmona, Carlos Alberto Reynoso, Carlos Aguilar, Nora E. Kerik, Luisa Rocha, Leticia Verdugo-Díaz, Víctor Treviño, José Bargas, Verónica Ramos-Mejía, Juan Fernández-Ruiz, Aurelio Campos-Romo, Iván Velasco

## Abstract

Human embryonic stem cells (hESCs) differentiate into specialized cells, including midbrain dopaminergic neurons (DAN). Non-human primates (NHPs) injected with 1-methyl-4-phenyl-1,2,3,6-tetrahydropyridine develop some alterations observed in Parkinson’s disease (PD) patients. We obtained DAN from hESCs and confirmed that they express dopaminergic markers, generate action potentials, and release dopamine (DA) *in vitro*. DAN were transplanted bilaterally in the putamen of parkinsonian NHPs. After grafting, animals improved motor behavior, evaluated by the HALLWAY task, and in agreement with this recovery, DA release was detected by microdialysis. Imaging techniques revealed changes in fractional anisotropy and mean diffusivity in magnetic resonance imaging and higher 11C-DTBZ binding in positron-emission tomography scans, associated with grafts. *Postmortem* analysis showed that transplanted DAN survived over ten months in the putamen, without developing tumors, in the immunosuppressed NHPs. These results indicate that cell replacement therapy with hESCs-derived DAN causes long-term biochemical, anatomical, and physiological improvements in this model of PD.

---

Parkinson’s disease (PD) is a neurodegenerative and progressive disorder caused by alterations in the basal ganglia circuitry after the degeneration of dopaminergic neurons (DAN) in the *substantia nigra pars compacta*, leading to a reduction of dopamine (DA) levels in the striatum. Clinically, PD is characterized by resting tremor, rigidity, and bradykinesia ^1,2^. The brain of non-human primates (NHPs) is anatomically and physiologically similar to the human brain. The mitochondrial complex I inhibitor 1-methyl-4-phenyl-1,2,3,6-tetrahydropyridine (MPTP) is a neurotoxin employed to develop NHPs PD models, because it reproduces some of its histological and motor alterations ^3–5^.

To provide therapeutic strategies, human pluripotent cells, such as embryonic stem cells (hESCs) and induced pluripotent stem cells, are used to generate DAN in vitro ^6–9^, by exposing them to molecules that stimulate signaling pathways found in ventral midbrain development. Mouse and human pluripotent stem cells-derived DAN, grafted into the striatum of 6-hydroxy-dopamine-lesioned rodents, exhibit long-term survival and improvements of rotation behavior and akinesia ^10–13^.

A protocol for differentiation of human pluripotent stem cells that produces homogeneous cultures of floor-plate derived mesencephalic DAN was used to graft MPTP-treated monkeys, and these neurons survived for a month without tumor formation ^12^. This method and others have been used to promote differentiation of induced pluripotent stem cells to DAN, which after grafting improved the behavioral alterations present in parkinsonian NHPs ^14–17^ and stabilized symptoms in one PD patient ^18^. Although behavioral recovery has been observed post-grafting in parkinsonian NHPs, there are no reports of DA release in the brain of transplanted animals.

Currently, imaging studies such as magnetic resonance imaging (MRI) and positron emission tomography (PET) allow the assessment of structural and functional changes in the brain. A long-lasting recovery on dopaminergic sites detected by PET has been reported after grafting DAN in the brain of NHPs ^14,16^, but changes detected in MRI post-grafting have not been analyzed. Here, we tested the therapeutic viability of differentiated DAN to recover the alterations present in parkinsonian NHPs. We used molecular, behavioral, biochemical and imaging techniques to comprehensibly study the impact of the graft, and found significant improvements associated with the presence of DAN in the putamen after ten months. This study reinforces the notion that cell replacement therapy might be effective in treating PD.

## Results

### Dopaminergic neuron differentiation

hESCs expressing enhanced Green fluorescent protein (EGFP) were employed ^19^. We first corroborated the expression of pluripotency markers on H9 wild-type hESCS by detecting *OCT4, SOX2, NANOG*, and *SSEA4* (Supplementary Fig. 1A). Subsequently, the H9-EGFP cell line was also confirmed to express *SOX2, OCT4*, and *KLF4* (Supplementary Fig. 1B), to present a euploid karyotype (Supplementary Fig. 1C) and to produced teratomas after inoculation in *nu/nu* immunodeficient mice (Supplementary Fig. 1D). Induction of midbrain floor-plate DAN was made as reported ^12^ in H9-EGFP cells. We confirmed the expression of relevant dopaminergic markers in agreement with a previous study ^12^, suggesting that our differentiation was successful: *LMX1A* and *FOXA2* were present from day 7 of differentiation and the expression of *Tyrosine Hydroxylase* (*TH*) was detected from day 21 (Fig.1A). RNA-Seq at different times of differentiation revealed that pluripotency genes were down-regulated as cell commitment proceeded with concomitant induction of DAN genes. These DA-related markers included, among others: *DDC, FOXA1, TH, LMX1A, HES6, PAX6, FOXA2, NR4A2, DCX, SYT4, NEUROG2, ASCL1* and *GLI3* (Fig. 1B, green arrows) at days 14 and 28. On the other hand, the pluripotency markers *NANOG, LIN28A, LIN28B, POU5F1* (Fig. 1B, red arrows) notably reduced its expression in these days, when compared to day zero. A comparison of the data of microarrays presented by Kriks and colleagues ^12^ with selected transcripts from our RNA-Seq results is presented in Supplementary Fig. 2. The full analysis of the sequencing data is presented elsewhere (Meléndez-Ramírez et al., submitted).

**Figure 1.**
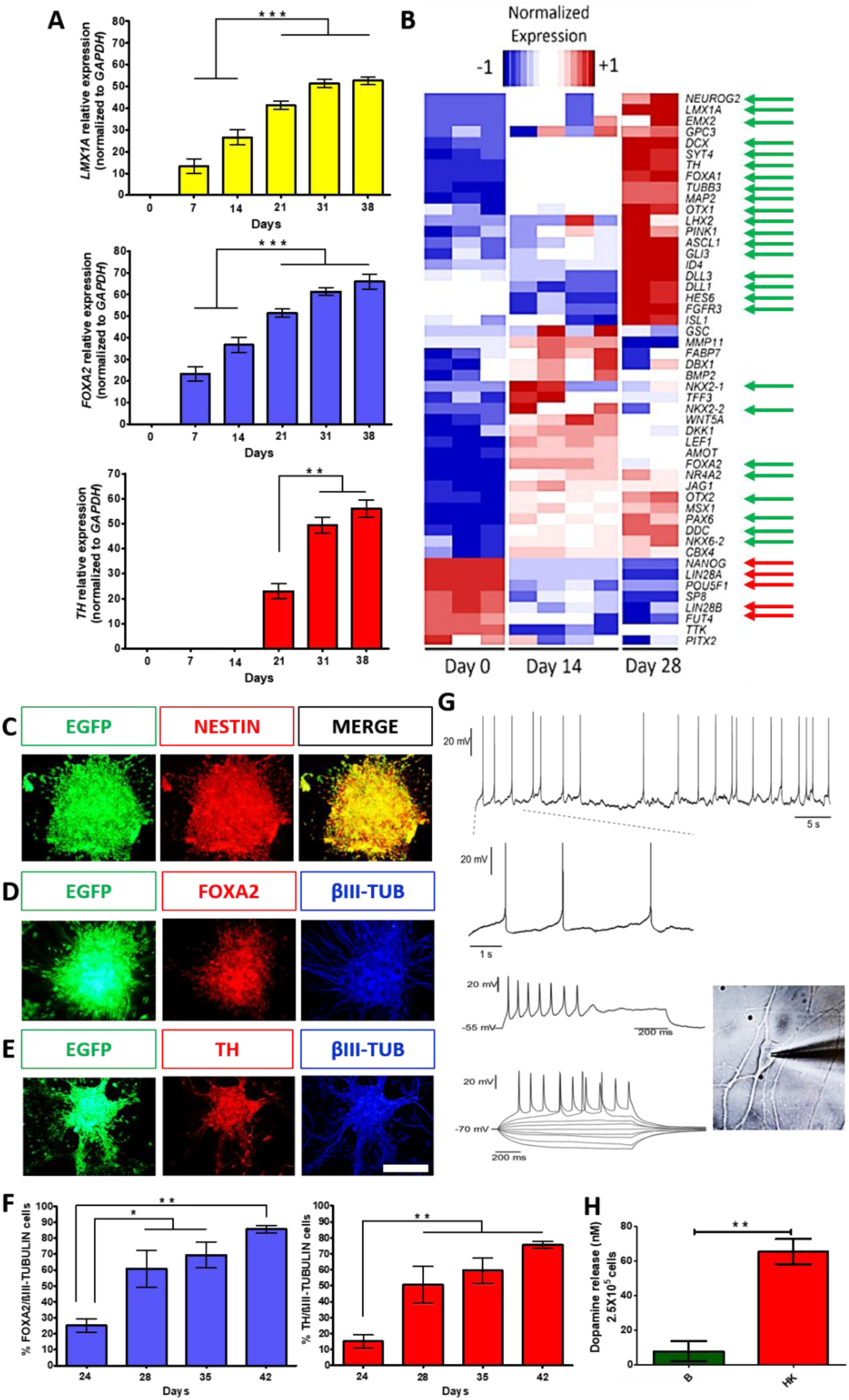
Dopaminergic differentiation of hESCs produces cells that have a transcriptional profile and electrophysiological properties characteristic of mature DAN, and release DA. **A)** Gene expression analysis, by RT-qPCR, of dopaminergic markers (*LMX1A, FOXA2* and *TH*) at different days of differentiation, normalized to *GAPDH*. Mean ± SEM; \**P*< 0.05, \*\**P* < 0.01, \*\*\**P* < 0.001, n=5 independent experiments. **B)** Heatmap from RNA-Seq data showing differential expression of genes at days 0, 14, and 28 of differentiation. Green arrows indicate significant upregulation of relevant DAN markers; red arrows point to significant downregulation of pluripotency markers. **C)** Immunocytochemistry at day 14 for expression of NESTIN and EGFP. **D)** Co-expression at day 35 for FOXA2 with βIII-TUBULIN (βIII-TUB). **E)** Co-expression at day 35 for TH with βIII-TUB. **F)** Quantitative co-expression analysis for FOXA2/βIII-TUBULIN (left panel) and TH/βIII-TUBULIN (right panel) at different days of differentiation. **G)** Electrophysiological recordings at day 60 shows spontaneous firing of action potentials and membrane potential oscillations at zero current (−54 mV of membrane potential), characteristic of DA neuron identity (1.5 Hz) (upper panel). Voltage responses to depolarizing and hyperpolarizing current injections (bottom panel) show evoked action potentials and depolarization block (middle panel) at high stimulus intensities; the phase contrast image of a patched neuron is shown on the right panel. **H)** DA levels in neurons at day 70 of differentiation in basal (B) solution and after chemical depolarization with isosmotic high-potassium (HK) medium. Mean ± SEM; \**P* < 0.05, \*\**P* < 0.01, \*\*\**P* < 0.001, n=5 independent experiments. Scale bar, 200 μm.

### Differentiated dopaminergic neurons have a mature phenotype

During the differentiation protocol, expression of the neural precursor marker NESTIN was confirmed on day 14 (Fig. 1C). At day 35, expression of the DAN markers FOXA2, TH, and βIII-TUBULIN was detected (Fig. 1D and E). Quantitative co-expression analysis for FOXA2/βIII-tubulin and TH/βIII-tubulin showed progressive and significant increases of double-positive cells, reaching 86% and 76% at day 42, respectively, (Fig. 1F). Electrophysiological recordings of differentiated cells between days 50 and 62 showed spontaneous single spike and bursting activity (6/10 neurons; Fig. 1G, top 2 rows) characteristic of DAN identity. Furthermore, all cells (n=10) elicited action potentials induced by current pulses (Fig. 1G, third and fourth rows). The neurons also showed rebound spikes in response to hyperpolarizing current pulses (n=6), rectification currents (n=8), and spontaneous synaptic activity (n=6) (data not shown). At day 70, cultures release DA, measured by HPLC, when stimulated with high potassium medium (Fig. 1H). Thus, this differentiation procedure generated mature DAN.

### hESC-derived dopaminergic neurons promote behavioral recovery in MPTP-treated NHPs

DAN produced from hESCs were grafted in parkinsonian NHPs, which develop motor alterations more similar to patients than rodent models. We performed imaging, behavioral and biochemical analyses (Fig. 2A). Animals were trained to perform the HALLWAY task before intoxication with MPTP and the effects of grafting hESCs-derived DAN were analyzed for 10 months, as previously described ^4,5^. To assess ambulatory and fine motor behavior, NHPs were filmed while walking through an acrylic hallway to reach the end, where 4 shelves, with two holes each, contained food rewards. Each NHP was trained to reach and eat two pellets on each shelf and return to the beginning of the hallway. Three successive behaviors, previously validated to be significantly modified after MPTP administration ^5^, were evaluated: displacement (time to cross the second third of the hallway); reaching (time to take the reward after passing the hand through the hole) and ingestion (time to introduce the reward into the mouth; Fig. 2B). The coordinates for grafting were obtained by MRI for each subject, in order to cover the entire putamen (Fig. 2C). MPTP administration significantly increased some task times, compared to the basal condition of NHPs(Supplemental videos). Sham grafting in one NHP did not induce overt behavioral improvement after ten months. In contrast, Grafted 1 had a significant decrease in performance times lasting ten months. Grafted 2 showed significant improvements compared to MPTP the first four months, with variable recovery afterward (Fig. 2D). These data demonstrate that bilateral DAN transplantation induces behavioral recovery.

**Figure 2.**
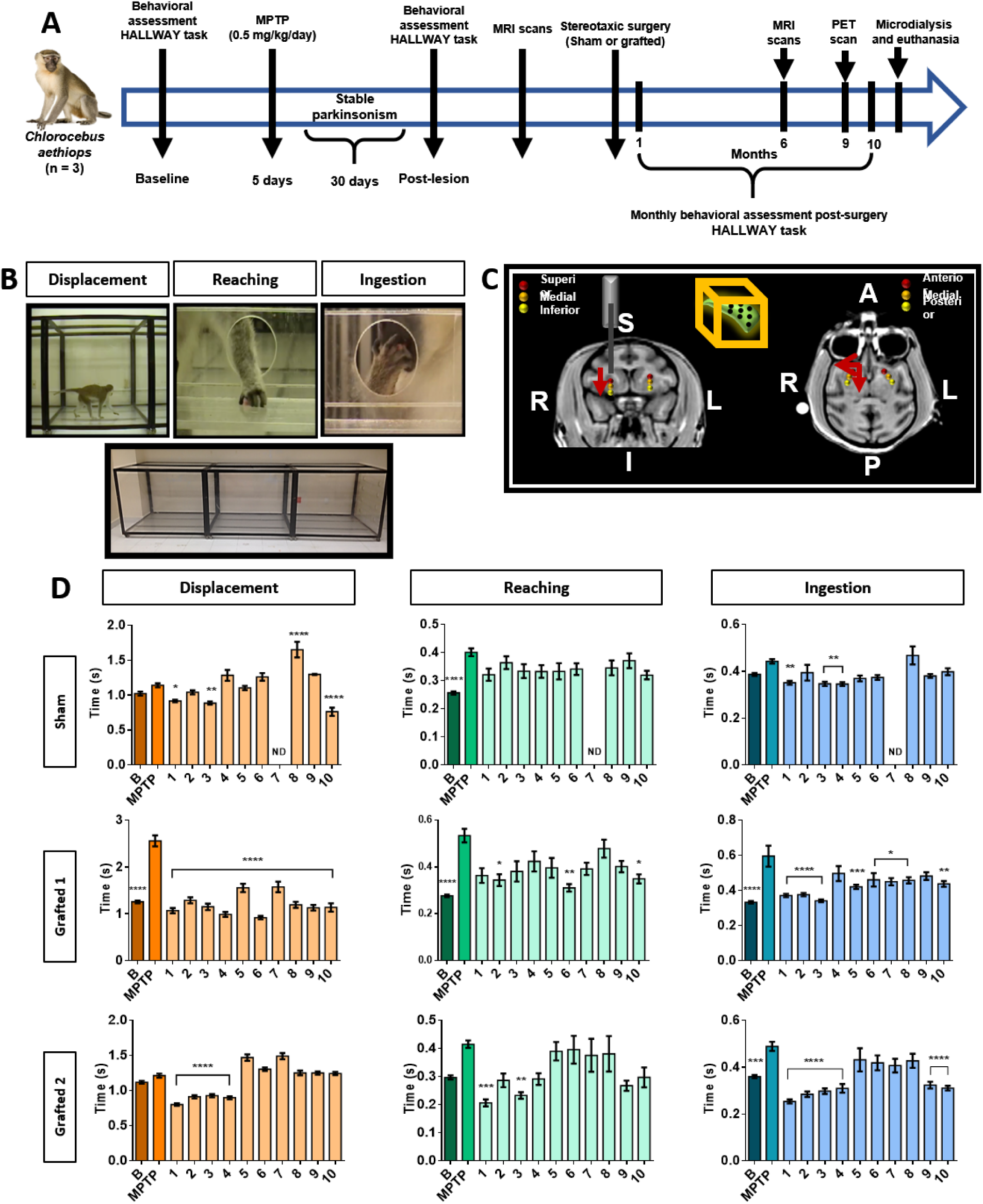
MPTP injection causes behavioral alterations in NHPs, that are diminished after transplantation of DAN. **A)** Timeline showing the experimental sequence for NHPs including behavioral assessment, MPTP administration, DAN grafting, PET and MRI scans, microdialysis and euthanasia. **B)** The panels show the design of the HALLWAY task and representative images of the assessed motor behaviors: displacement, reaching and ingestion of the reward. **C)** MRI images depict the bilateral grafting sites for DAN into the putamen. Red arrows indicate the direction of each microinjection (R: right; L: left; S: superior; I: inferior; A: anterior; P: posterior). The box in the middle top part represents the 9 cell deposits in a putamen. **D)** Graphs show the average time it took each NHP to perform each motor task. B, basal; MPTP, lesioned; 1-10, month evaluated; N.D., Not determined. Mean ± SEM; \**P* < 0.05, \*\**P* < 0.01, \*\*\**P* < 0.001, \*\*\*\**P* < 0.0001 relative to the MPTP stage.

### Diffusion tensor imaging

To assess cellular and axonal density changes by MRI, we performed fractional anisotropy (FA) and mean diffusivity (MD) analyses, respectively. Before grafting, we analyzed the overall changes in the left and right putamen of healthy and MPTP-injected NHPs, as depicted in Supplementary Fig. 3A. FA decreased in MPTP-treated NHPs when compared to a healthy group, showing reductions of 6.6 % in the right putamen and 5.4 % in the left putamen (Supplementary Fig. 3B). The MD comparison between healthy and MPTP-treated NHPs showed an increase of 11.4 % in the right putamen and 5.8 % in the left putamen (Supplementary Fig. 3B).

We then compared FA and MD measured after MPTP treatment with the post-operative condition (POp: Sham or Grafted). The striatum is a heterogeneous structure that includes anatomic/functional subdivisions and several models have been proposed ^20,21^. We decided to use the anatomo-functional subdivisions that designate the anterior or limbic putamen as involved in motivation, the medial or associative putamen related to cognition and the sensorimotor or posterior putamen, linked to locomotion ^22^ and defined these 3 regions for further analysis at 6 months post-surgery (Fig. 3A).

**Figure 3.**
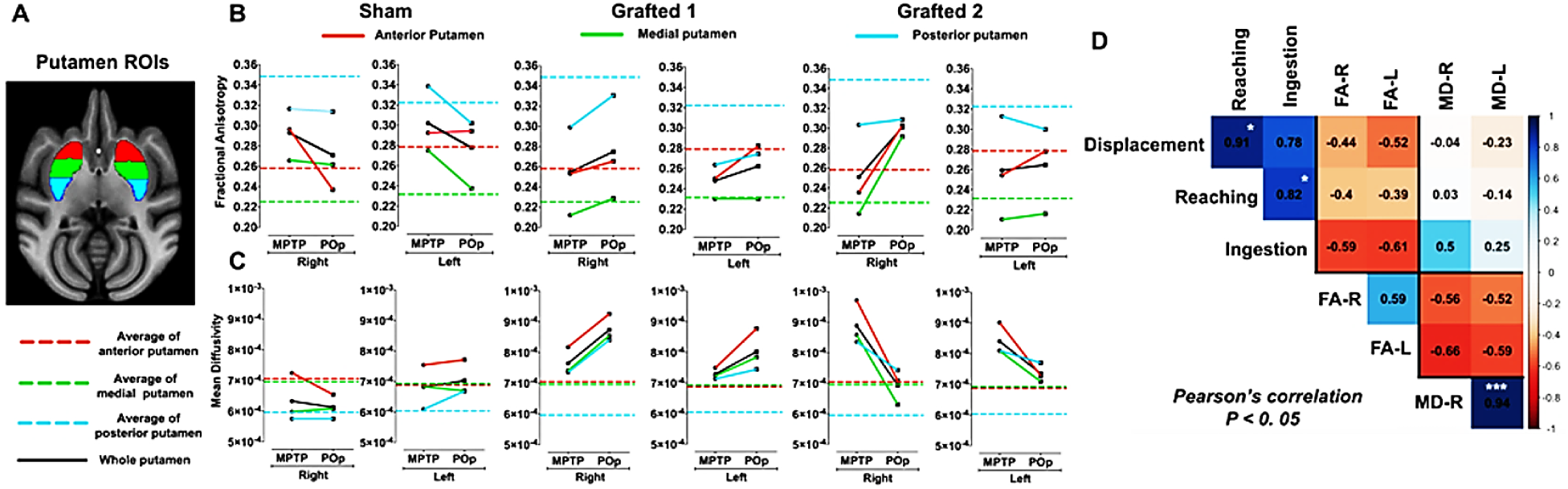
FA and MD measurements in MPTP-treated NHPs after sham surgery or DAN transplantation. **A)** MRI template and ROIs of putamen: anterior (red), medial (green) and posterior (blue). **B)** FA and MD **(C)** quantification in Sham and Grafted NHPs in bilateral putamen in MPTP condition and post-operative (POp), respectively. Reference dotted lines represent the average FA and MD of anterior (red), medial (green) and posterior (blue) putamen in the healthy group. The black lines represent the average of FA and MD of anterior, medial and posterior putamen. **D)** Pearson’s correlation matrix between the unilateral FA and MD of whole putamen with the behavioral scores (displacement, reaching and ingestion) of all NHPs in both MPTP and POp conditions. \**P* < 0.05, \*\*\**P* < 0.005.

In the Sham NHP, FA measures decreased after sham surgery, with exception of the left anterior putamen. In sharp contrast, FA increased bilaterally in Grafted 1 and Grafted 2 animals, compared to their previous MPTP condition, except for the posterior left side of Grafted 2; when considering the 3 portions, Sham showed a decrease and Grafted NHPs an increase post-operatively (POp) in FA, illustrated by the black lines (Fig. 3B). On the other hand, MD values in Sham NHP rendered a variable response after surgery, although the changes were very discreet. Interestingly, Grafted 1 had increases in all the analyzed regions, and Grafted 2 showed a consistent bilateral reduction post-grafting (Fig. 3C). To correlate behavioral changes with DTI measures in all NHPs in both MPTP and POp conditions, a Pearson’s correlation analysis was performed (Fig. 3D). The resulting matrix shows a positive and significant correlation between reaching with both displacement and ingestion. Also, measures of FA in both hemispheres present a positive correlation of 0.59. For MD, the correlation between the left and right sides was positive and highly significant. When comparing behavioral tests with FA, values were between -0.61 and -0.39, showing a negative correlation with displacement, reaching, and ingestion, which indicates increased values of FA and decreased times to perform motor tasks. In contrast, MD of right and left putamen have values close to zero for displacement and reaching, and a positive non-significant correlation with ingestion. Interestingly, the correlation between FA and MD was negative, with values from -0.66 to -0.52.

### PET qualitative analysis

We performed ^11^C-DTBZ PET scanning to assess the presence of DA nerve terminals, due to an incomplete lesion, or to detect somata of grafted DAN, in the putamen. Sham NHP showed residual ^11^C-DTBZ binding, but its z-score is lower than Grafted 1 and Grafted 2 (Fig. 4). Grafted 1 had greater binding at anterior regions, compared to Sham; Grafted 2 presented higher signal at posterior levels. These data revealed that Grafted NHPs, at 9 months post-surgery, have increased ^11^C-DTBZ binding in both putamens, indicative of functional DAN at the transplanted sites.

**Figure 4.**
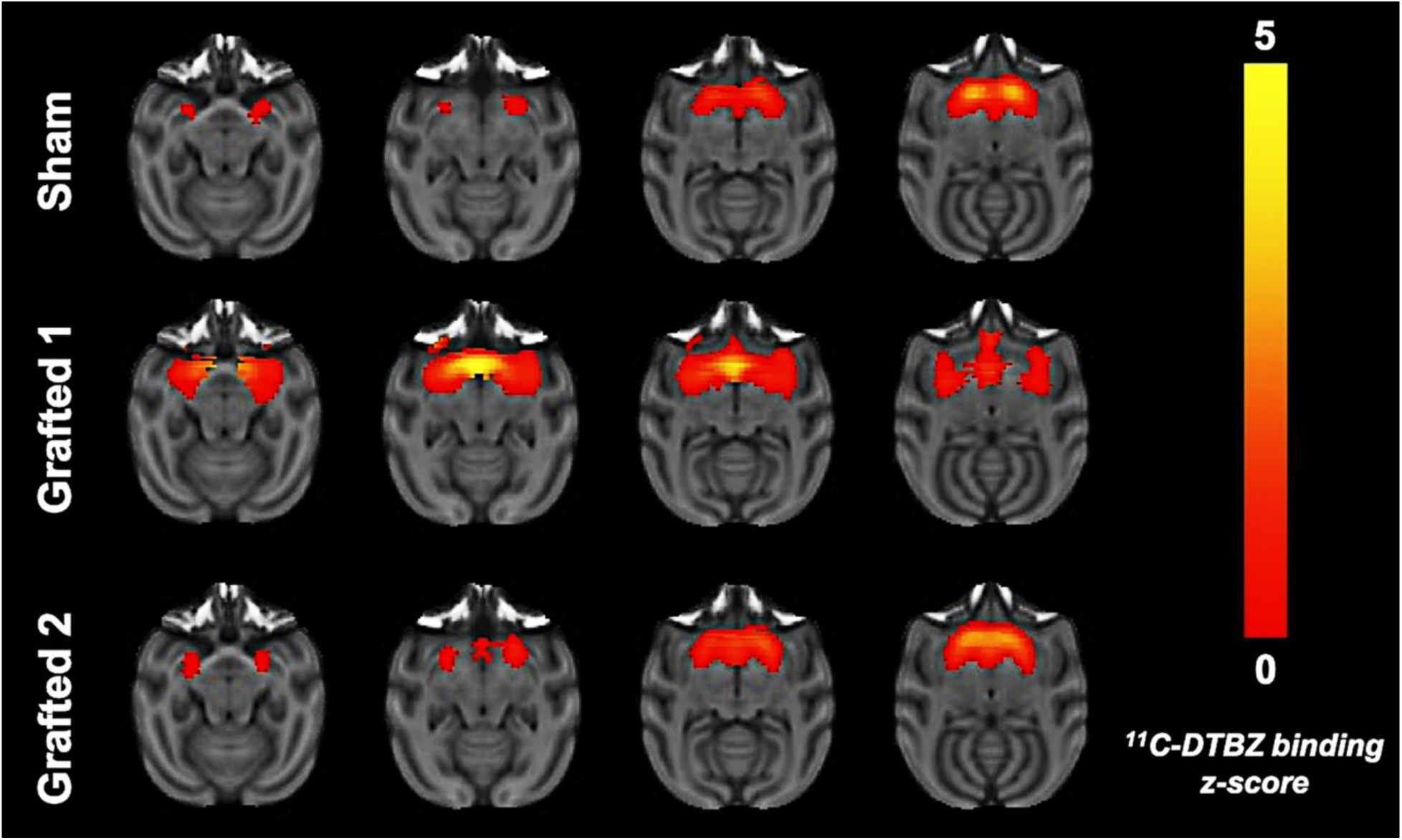
Axial PET sections 9 months after transplantation, illustrating a higher binding potential of 11C-DTBZ in the brains of Grafted NHPs. ^11^C-DTBZ binding was measured individually for Sham, Grafted 1 and Grafted 2. All PET images are merged onto MRI-T1 acquisitions. The warm color bar represents the normalized *z-score*.

### Transplanted neurons release dopamine

Before euthanasia, microdialysis was used to quantify the extracellular DA, released in the putamen after chemical stimulation *in vivo*. Probes were inserted simultaneously in both hemispheres. Sham NHP had low levels of basal DA and did not present a clear response after stimulation with high potassium or amphetamine, administered through the dialysis membrane. In both Grafted NHPs, baseline DA concentrations were elevated. Notably, extracellular DA concentrations sharply increased in both transplanted sides after both stimuli (Fig. 5). The metabolite DOPAC decreased when administering the solution with high potassium and amphetamine in Grafted NHPs (Fig. 5). These data demonstrate that DA is released at the sites of grafting.

**Figure 5.**
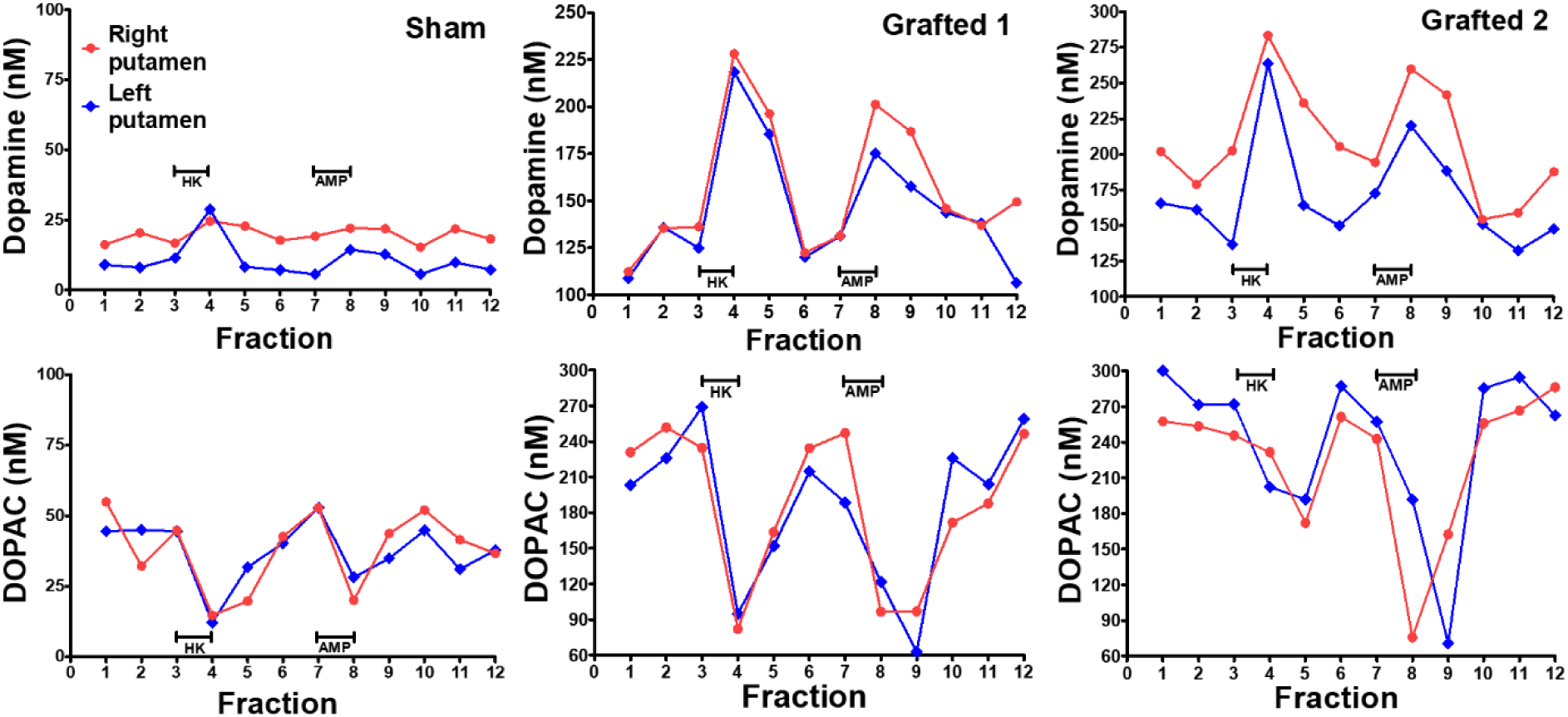
Dopamine is released in the putamen, after chemical stimulation, in Grafted NHPs ten months post-surgery. Time-course on DA and DOPAC concentrations measured by microdialysis in each putamen after stimulation with 100 mM KCl isosmotic medium (HK) or 30 μM amphetamine (AMP) in Sham, Grafted 1 and Grafted 2 NHPs. Each fraction was collected for 12:30 minutes. DOPAC, 3,4-dihydroxyphenylacetic acid.

### Dopamine neurons survive for ten months in the putamen of MPTP-treated NHPs

After fixation, brain slices were analyzed for the presence of grafted GFP+ neurons in each putamen (Fig. 6A). These DAN were TH+, GIRK2+, and MAP2+ (Fig. 6B and 6D). Importantly, grafted animals did not develop tumors. TH counting showed 2.5×10^5^ and 2×10^5^ cells per hemisphere in Grafted 1 and 2, respectively. Of note, MPTP injection did not cause a complete loss of DAN cells in the *substantia nigra*, since all animals had 5×10^4^ TH+ cells per hemisphere (Fig. 6 C).

**Figure 6.**
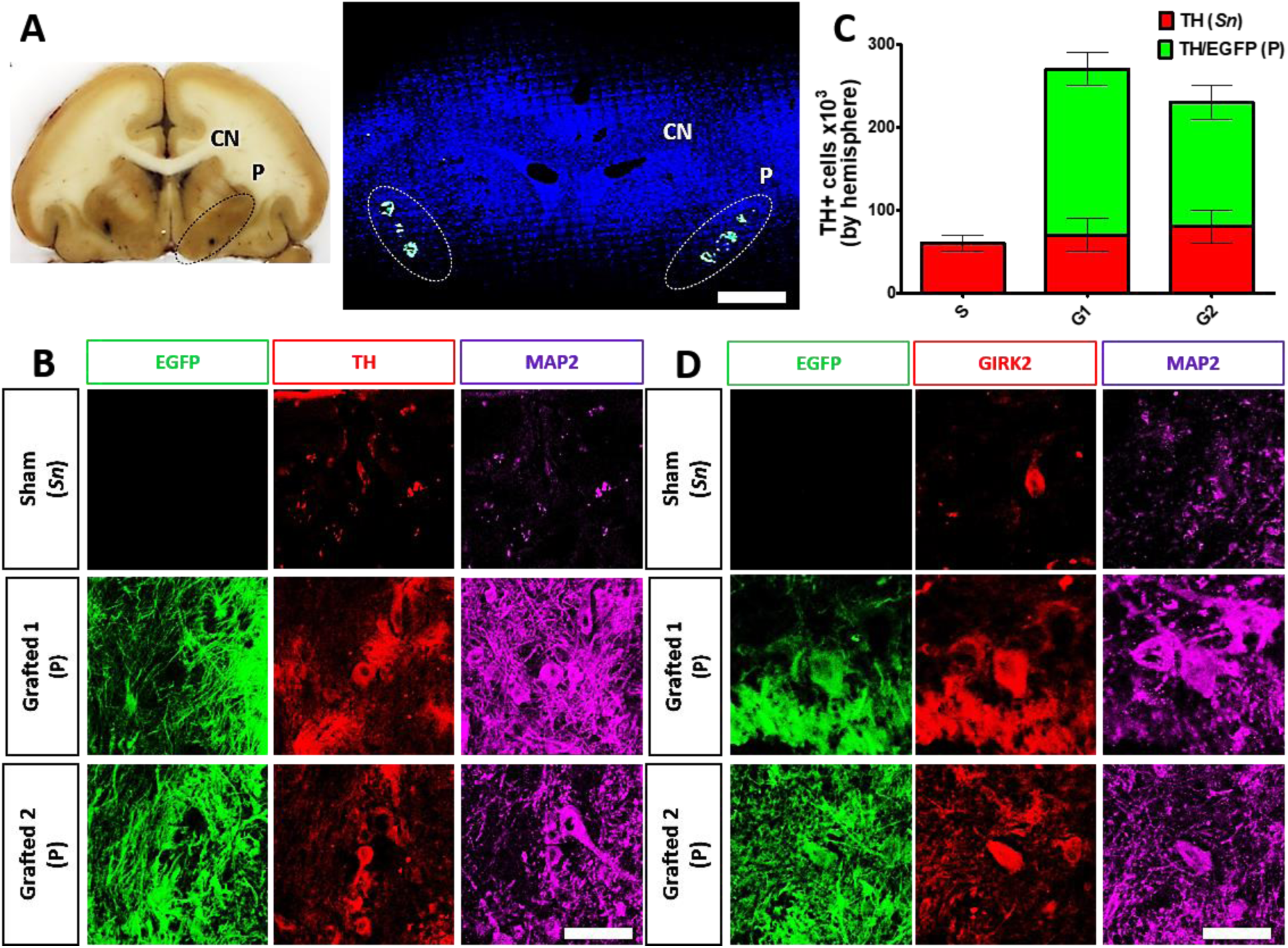
Survival of human DAN grafted in the putamen after 10 months. **A)** Representative coronal section of the brain showing the caudate nucleus (CN) and the putamen (P) (left panel). On the right side, a confocal reconstruction of a coronal section is presented after nuclear staining (blue) to show the localization of GFP+ transplant sites at each putamen (dotted lines) in Grafted 1. Scale bar, 1 cm. **B)** Surviving cells in the *substantia nigra* (*Sn*) and co-expression of TH (red), MAP2 (cyan) and EGFP (green) by grafted DAN in the putamen (P) after ten months. **C)** Quantitative analysis for endogenous TH+ (*Sn*) and TH+/EGFP+ (P) cells after grafting. Mean ± SEM. **D)** Co-expression for GIRK2 (red), MAP2 (cyan) and EGFP (green). Scale bar for B and D, 200 μm.

## Discussion

Human DAN transplanted in the putamen of parkinsonian NHPs improved motor behavior, increased ^11^C-DTBZ binding, and survived for ten months without tumors. Importantly, grafted human neurons released DA in the brain and generated changes in FA and MD, which to our knowledge, is the first time reported in parkinsonian NHPs.

The derivation of midbrain DAN from hESCs was efficient, due to the neural-inducing properties of specific small molecules and supported with an 76-86% of DAN that co-expressed FOXA2/βIII-tubulin and TH/βIII-tubulin at day 42 of differentiation. Other data that corroborate the identity of these neurons were obtained by RNA-Seq and RT-qPCR, showing the expression of relevant specific markers (*FOXA1, FOXA2, TH, LMX1A, SYT4, DDC, PAX6, DCX, NEUROG2, ASCL1*). The full maturation of these neurons was at 50 days of culture, corroborated by electrophysiology and the release of DA to the medium, which is consistent with previous reports ^12,16,23,24^.

Previous reports of grafting DAN differentiated from human stem cells showed that cell therapy induce improvements in parkinsonian NHPs when motor behavior and DA uptake sites were assessed by PET ^16,17,25^. Among the available behavioral tests, we found that the HALLWAY task is not just sensitive to MPTP lesion, but also a good approach to evaluate behavior recovery after cell replacement therapy ^4,5^. It remains difficult to have a behavioral test with low variability ^26^, so we decided to correlate the motor improvement over a period of ten months, with DA release and imaging analyses. Although the Sham NHP had some months in which the displacement and ingestion were significantly lower than MPTP, this might be due to the plasticity of MPTP-treated brain ^27^. Nonetheless, the recovery observed in both Grafted NHPs was clear and correlated with changes in MRI and *in sit*u DA release. This study reports a long-term recovery after grafting, although the number of animals used was small, we found a good correlation within the different approaches that we performed.

The anisotropy of water diffusion in brain tissue is modified by development or disease and can be evaluated by FA and MD derived from DTI. In PD, there is a reduction of FA in basal ganglia nuclei ^28^ similar to our observations in the putamen of all NHPs after MPTP administration, suggesting that FA decreases reflect microstructural damage of tissue integrity ^29–31^. MD increases are also related to tissue loss and extracellular fluid accumulation, which we also detected after MPTP administration. The decrease of FA and the increase of MD might be due to the loss of dopaminergic terminals into the putamen nuclei and the retraction of striatal medium spiny neurons after MPTP denervation ^32–34^. The functional recovery promoted by cell transplantation was accompanied by higher FA values in our longitudinal comparison along the putamen. Interestingly, FA values gradually increase, and MD values gradually decrease due to the axonal growth, ion flux across axons, gliosis, and glial scar formation after spinal cord injury in rats ^35,36^. Although in our study, both Grafted NHPs showed an increase of FA, Grafted 1 showed an increment of MD whereas Grafted 2 showed a reduction of MD. Our correlation analysis of DTI results revealed a negative correlation between FA and behavioral performance, but also between FA and MD. Such changes, especially increased FA, should be useful to monitor the survival and functioning of DAN *in vivo* with a non-invasive, non-radioactive technique.

Microdialysis had not been used to directly measure DA levels in grafted NHPs. Ten months after grafting, and before euthanasia, grafted neurons released DA with specific stimuli *in vivo*, similar to findings reported in rodent models of PD ^37,38^. Both Grafted NHPs showed higher levels of DA, compared to Sham, which partially explains sustained behavioral recovery. Further studies are required to establish whether postsynaptic DA receptors do not present supersensitivity after grafting NHPs, a phenomenon that has been reported after DAN transplantation in hemiparkinsonian rats ^37^.

PET analysis showed that grafted DAN were functional *in vivo*, suggesting the presence of DAN into the brain ^39^, which may be associated with favorable changes in the behavior. MPTP administration to NHPs reduces the binding of dopaminergic PET probes ^26,40^. Consistent with our results, DAN grafting to parkinsonian NHPs consistently increased PET signals and this has been correlated to behavioral improvement ^16,41^.

A concern about hESC-based therapies is the development of tumors, considering that few pluripotent cells can remain undifferentiated ^42^. Ten months after cell transplantation NHPs did not show any tumors. Additionally, RNA-seq showed that the pluripotency genes, as well as the cell proliferation gene *TTK*, decreased upon differentiation, which might partially explain the lack of tumors in Grafted NHPs ^43^. Three percent of grafted DAN survived in the long-term, similar to previous reports ^16,17^, showing that a high proportion of grafted neurons die shortly after surgery. Despite this, DAN survival can form new tracts and generate behavioral improvements. These neurons still express TH and partially compensate for the loss of DA. According to the development of hPSC-derived DAN replacement therapy in PD, the survival of a minimal number of grafted DAN and enough reinnervation in the putamen is necessary for the improvement of symptoms ^13^. These data suggest that an improvement in motor function is achieved when a proper number of grafted DAN survive. This study supports the notion that stem cells differentiate to DAN might be grafted into PD patients.

## Materials and methods

### Dopaminergic differentiation of hESCs

Human embryonic stem cell line H9 (WA09)-GFP, which constitutively express enhanced Green Fluorescent Protein (EGPF), was used for all experiments ^19^. Prior to differentiation, cells were expanded in supplemented Knock Out (Gibco ®) medium, conditioned by mitotically inactivated mouse embryonic fibroblasts (MEFs), with fresh FGF-2 (10 ng/ml; Sigma), over a Matrigel (BD ®) matrix, until they reached 75% confluency. The floor-plate dopaminergic differentiation protocol ^12^ was performed with minor modifications ^44^. Pluripotent hESCs were incubated with small molecules to start the dual SMAD inhibition: BMP and TGF-b receptors were pharmacologically blocked with 100 nM LDN193189 (Stemgent) and 10 mM SB431542 (Tocris), respectively. Wnt canonical pathway was stimulated by inhibiting the kinase GSK-3b, with 3 mM CHIR99021 (Stemgent). Shh signaling was stimulated by 1 mM SAG (Sigma) and 2 mM purmorphamine (Stemgent). Recombinant human FGF-8 was added at 100 ng/ml (Peprotech) until day 7. At day 14, cells were cultured in Neurobasal medium with B27 supplement and the morphological changes were evident; cells were positive in a high proportion to NESTIN, indicating that neural progenitors were present at this stage. Neuronal differentiation and survival were promoted by 20 ng/ml BDNF (Peprotech), 0.2 mM ascorbic acid (Sigma), 20 ng/ml GDNF (Peprotech), 1 ng/ml TGF-β3 (Peprotech), 0.5 mM dibutyryl cAMP (Sigma) and the Notch inhibitor DAPT at 10 mM (Sigma) in the culture medium. On day 22, cells were dissociated using TrypLE Express (Life Technologies) and either seeded onto poly-l-ornithine, Fibronectin- and Laminin-treated plates or used for grafting. For electrophysiological and HPLC assays, cultures were grown for over 50 days of differentiation in NDM.

### RT-qPCR

RNA was isolated using TRIzol Reagent (Thermo Fisher Scientific). Complementary DNA (cDNA) was synthesized by SuperScript III Reverse Transcriptase (Thermo Fisher Scientific) and used for RT-PCR amplification (Taq DNA Polymerase, Thermo Fisher Scientific). Amplification of 50 ng of cDNA was performed with the QuantiFast SYBR Green PCR Master Mix (Qiagen) with a StepOnePlus RealTime PCR System (Applied Biosystems) with the following primers:

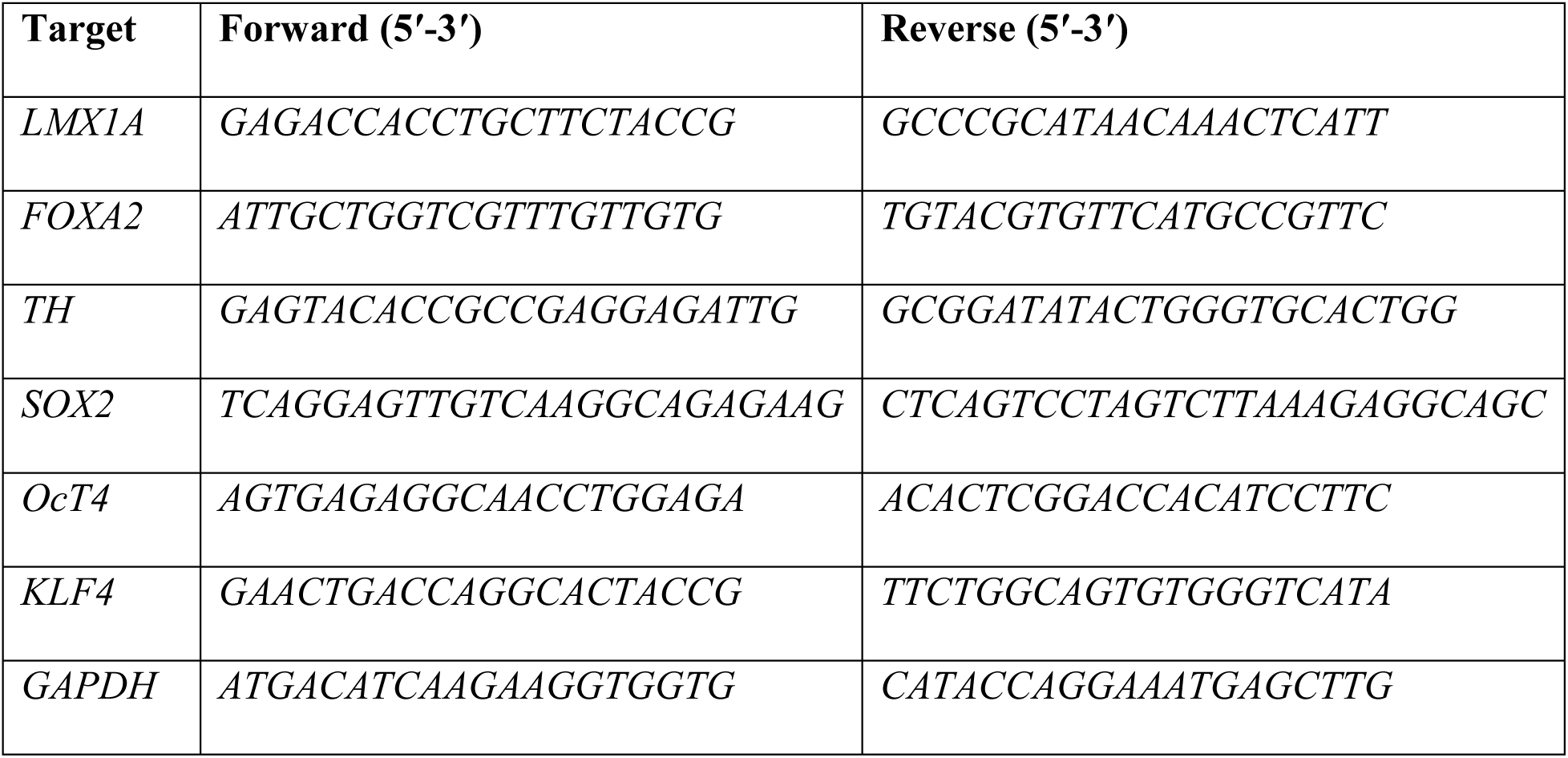

### Immunocytochemistry

Cells were fixed with 4% paraformaldehyde, permeabilized and blocked with 0.3% TritonX-100 and 10% normal goat serum in PBS, and incubated overnight with the primary antibodies in PBS plus 10% normal serum:

| Antibody | Host | Dilution | Brand | Catalog Number |
| --- | --- | --- | --- | --- |
| Oct4 | Mouse | 1:250 | BD Biosciences | 611202 |
| Sox2 | Rabbit | 1:500 | Abcam | AB97959 |
| Nanog | Rabbit | 1:1000 | Peprotech | 500-P236 |
| SSEA4 | Mouse | 1:400 | Abcam | AB16287 |
| NESTIN | Rabbit | 1:500 | Covance | 839801 |
| TH | Rabbit | 1:500 IC<br>1:1000 IH | Pel-Freez Biologicals | P40101 |
| βIII-TUBULIN | Mouse | 1:1000 | Covance | MMS435P |
| HNF3β/FoxA2 | Rabbit | 1:500 | Millipore | 07633 |
| MAP2 | Mouse | 1:500 | SIGMA | M4403 |
| GIRK2 | Mouse | 1:1000 | Millipore | AB5200 |
| Alexa Fluor 350<br>Anti-Mouse<br>IgG | Goat | 1:1000 | Thermo Fisher<br>Scientific | A21049 |
| Alexa Fluor 488<br>Anti-Mouse<br>IgG | Goat | 1:1000 | Thermo Fisher<br>Scientific | A11029 |
| Alexa Fluor 568<br>Anti-Rabbit<br>IgG | Goat | 1:1000 | Thermo Fisher<br>Scientific | A11036 |

Negative controls without primary antibodies were included and showed no unspecific staining. A Nikon eclipse TE2000-U microscope was used for image acquisition and the Image-Pro Plus program Version 4.5 (MediaCybernetics, Bethesda, MD) was employed for epifluorescence image analysis.

### Karyotype analysis

Chromosomal analysis was performed by GTG-banding analysis at Reproducción y Genética AGN, Hospital Ángeles del Pedregal, México. Briefly, cells were incubated with colcemid (2 h), harvested by trypsinization, processed with hypotonic solution, and fixed with methanol:acetic acid (3:1). Metaphases were spread on slides and chromosomes were counted and classified using the G banding technique.

### Teratoma formation assay

All mouse procedures were performed in accordance with current Mexican legislation (NOM-062-ZOO-1999, SAGARPA), the Guide for the Care and Use of Laboratory Animals of the National Institutes of Health (NIH) and approved by the institutional Animal Care and Use Committee of IFC-UNAM. hESC were grown on Matrigel (BD Biosciences), harvested by trypsinization, washed in PBS and re-suspended in KSR medium with 30% of Matrigel. Cells from a T-25 flask at 80% confluence containing approximately 3-5×10^6^ were subcutaneously inoculated into the dorsal flank of each 6–8-week-old nude (nu/nu) mice. The presence of cells from the three embryonic germ layers was determined by hematoxylin and eosin-stained sections.

### RNA-Seq

Massive next generation sequencing was performed on a MiSeq equipment from illumina by paired end reads (2×75) using TruSeq RNA Stranded mRNA Library Prep following the provider protocol. About 50.4 million passed-filter reads were obtained. On average, 10.98% reads per sample were obtained (SD=1.95%). Of these, 3 replicates represented day 0 (11.81%, 13.30% and 11.62%), four represented day 14 (10.48%, 11.24%, 10.99% and 12.65%), and two represented day 28 (6.45% and 10.30%) of differentiation. Similar gene expression patterns were found when we compared our results with the gene expression levels previously reported by microarrays ^12^. The standardized Z-Score was used to illustrate the expression patterns.

### Electrophysiological analysis

Whole-cell patch-clamp recordings were performed on 50- to 65-days-old cultures of hESCs differentiated into DAN. Neurons were transferred to a recording chamber and continuously superfused with oxygenated saline solution (4-5 ml/min) at room temperature (∼25°C). Micropipettes were pulled (Sutter Instrument, Novato, CA) from borosilicate glass tubes (WPI, Sarasota, FL) to an outer diameter of 1.5-mm for a final DC resistance of 4–6 MΩ when filled with internal saline containing (in mM): 120 KSO3CH4, 10 NaCl, 10 EGTA, 10 HEPES, 0.5 CaCl2, 2 MgCl2, 2 ATP-Mg, and 0.3 GTP-Na (pH = 7.3, 290 mOsm/l). Neurons were visualized with infrared differential interference video microscopy and epifluorescent illumination with a 40X immersion objective (0.8 NA; Nikon Instruments, Melville, NY) and a CCD camera (Cool Snap ES2, Photometrics, Tucson AZ, USA). Recordings were made with an Axopatch 200A amplifier (Axon Instruments, Foster City, CA) and data were acquired with Im-Patch© an open source software designed in the LabView environment (National Instruments, Mexico City, Mexico; available at www.im-patch.com). The giga-seal resistances were in the range of 10–20 GΩ. The current signals from the amplifier were filtered at 5 kHz through a four-pole low-pass filter.

### Lesion of non-human primates with MPTP

Experiments were performed in three adult Vervet monkeys (*Chlorocebus aethiops*), 2 females, and 1 male, aged 10-15 years and weighing 4.0-4.5 kg. NHPs were housed in individual cages with 12 h light/dark cycle, room temperature at 23 ± 1 °C, 50 ± 10 % of relative humidity and were fed twice daily with a diet of High protein monkeys LabDiet 5038 biscuits (High Protein Monkey Chow of Lab Chows, Purina®), water *ad libitum*, and fresh fruits and vegetables. All procedures were performed in accordance with current Mexican legislation NOM-062-ZOO-1999 (SAGARPA), the Guide for the Care and Use of Laboratory Animals of the National Institutes of Health (NIH) and approved by the Animal Care and Use Committees of Instituto Nacional de Neurología y Neurocirugía (20/13) and Universidad Nacional Autónoma de México. An additional group of three healthy NHPs, 2 females and 1 male, were imaged with MRI to study the effect of MPTP.

Bilateral parkinsonism was induced in the 3 NHPs by intramuscular MPTP hydrochloride (Sigma-Aldrich, St Louis, USA) administration, dissolved in saline at 0.5 mg/kg to reach 2-2.5 mg/kg divided into 4 or 5 daily injections until they developed an extra-pyramidal syndrome ^4,5^. MPTP administration was made in a designated special room with the necessary safety measures for the animals and the personnel, following security protocols ^45^. After MPTP administration, NHPs were closely monitored by a veterinary and were provided with water, food pellets and fresh fruits to maintain their corporal weight and general well-being.

### Motor behavioral assessment

Parkinsonian motor symptoms were rated using a HALLWAY task previously reported to evaluate the free movements of NHPs ^4,5^. Ten-minute sessions were video recorded per day for five consecutive days. Two observers blind to treatment scored motor performance independently for each monkey. The analysis of the videos consisted in frame-by-frame quantifications of three motor behavioral parameters: 1) Displacement: time to cross the second third of the hallway; 2) Reaching: time to take the rewards and 3) Ingestion: time to bring the reward to the mouth. NHPs were first recorded for five days before MPTP administration. After MPTP intoxication, NHPs had a one-month parkinsonism stabilization period, before performing again the behavioral task for five days. After sham or grafting surgery, the animals were evaluated for five days monthly over a period of 300 days (Supplementary videos).

### Magnetic resonance imaging

MRI images were acquired in a GE Discovery MR750 (General Electric Healthcare, Milwaukee, WI) scanner of 3T and a commercial 32-channel head coil array at the Instituto de Neurobiología of Universidad Nacional Autónoma de México. The animals were anesthetized with Tiletamine/Zolazepam (125/125 mg) at a dose of 2 mg/kg (i.m.). T1-weighted images were acquired by Fast Spoiled Gradient Echo sequence using the following parameters: TR/TE = 8.5/3.2 ms; FOV = 256 × 256 mm^2^; reconstruction matrix = 256 × 256; final resolution = 1×1×0.5 mm^3^. The diffusion weighted imaging (DWI) were acquired by Single Shot Echo Planar Imaging sequence with the following values: TR/TE = 6500/99 ms; FOV = 256 × 256 mm^2^; reconstruction matrix = 256 × 256; final resolution = 1×1×2 mm; 35 slices; slice thickness = 2 mm; volumes = 65; no diffusion sensitization images with b = 0 s/mm^2^ and 60 DWI of independent directions with b = 2000 s/mm^2^.

### Surgical procedure and DAN transplantation

The surgical procedure was performed 6 months after MPTP administration. Lesioned NHPs underwent stereotactic surgery. The surgical coordinates were calculated for each subject from T1 images by MRI as follows:

|  |  | (Sham) |  |  |  |  |  | (Grafted 1) |  |  |  |  |  | (Grafted 2) |  |  |  |  |  |
| --- | --- | --- | --- | --- | --- | --- | --- | --- | --- | --- | --- | --- | --- | --- | --- | --- | --- | --- | --- |
| Graft |  | Left<br>(cm) |  |  | Right<br>(cm) |  |  | Left<br>(cm) |  |  | Right<br>(cm) |  |  | Left<br>(cm) |  |  | Right<br>(cm) |  |  |
| A | L | 4.50 |  |  | 6.35 |  |  | 4.68 |  |  | 6.58 |  |  | 4.26 |  |  | 6.46 |  |  |
|  | A<br>P | 4.90 |  |  | 4.90 |  |  | 5.1 |  |  | 5.1 |  |  | 4.2 |  |  | 4.2 |  |  |
|  | V | 3.21 | 3.01 | 2.82 | 3.41 | 3.21 | 3.02 | 3.06 | 2.76 | 2.46 | 3.06 | 2.76 | 2.46 | 3.22 | 2.92 | 2.72 | 3.22 | 2.92 | 2.72 |
| M | L | 4.35 |  |  | 6.50 |  |  | 4.58 |  |  | 6.68 |  |  | 4.16 |  |  | 6.56 |  |  |
|  | A<br>P | 4.70 |  |  | 4.70 |  |  | 4.9 |  |  | 4.9 |  |  | 3.9 |  |  | 3.9 |  |  |
|  | V | 3.21 | 3.01 | 2.82 | 3.41 | 3.21 | 3.02 | 3.06 | 2.86 | 2.66 | 3.06 | 2.86 | 2.66 | 3.22 | 2.92 | 2.72 | 3.22 | 2.92 | 2.72 |
| P | L | 4.20 |  |  | 6.65 |  |  | 4.48 |  |  | 6.78 |  |  | 4.06 |  |  | 6.76 |  |  |
|  | A<br>P | 4.41 |  |  | 4.41 |  |  | 4.7 |  |  | 4.7 |  |  | 3.7 |  |  | 3.7 |  |  |
|  | V | 3.21 | 3.11 | 3.01 | 3.41 | 3.31 | 3.21 | 3.06 | 2.86 | 2.66 | 3.06 | 2.86 | 2.66 | 3.12 | 2.82 | 2.62 | 3.12 | 2.82 | 2.62 |

The stereotaxic zero was placed in the middle line at auditory canal levels; from there, we calculated the distance to the target area. Two monkeys received DAN (designated Grafted 1 and Grafted 2) and one monkey received culture medium (Sham). All animals were fasted overnight before surgery. The induction of anesthesia was performed with Tiletamine/Zolazepam (125/125 mg) at a dose of 2 mg/kg of weight (i.m.). Then, NHPs were intubated and anesthetized with isoflurane (1-2 %, O_2_ flow rate of 2 l/min) to maintain a proper anesthetic state. NHPs body’s temperature was maintained by a heating blanket and its head was placed onto the stereotaxic frame. Target areas were shaved, sterilized and bilateral incisions were made on the scalp to expose the cranial surface. The skull was drilled making vertical holes to inject the cells into each putamen. 22-day cultures were dissociated, and cell re-suspended at 8.8 × 10^5^ cells/µl. Grafts of DAN were distributed in nine deposits at three different antero-posterior sites totaling 8 × 10^6^ cells per hemisphere. In each injection site, cells were slowly delivered in three different locations in the dorsoventral axis. After injection, the burr-hole was sealed with bone wax and the fascia, muscle and skin were sutured. Post-operative care started at the end of the surgery and followed for 1-week with analgesic (tramadol 1.4 mg/kg, i.m.) and antibiotic (cephalexin: 25 mg/kg orally). All NHPs received immunosuppression by oral cyclosporine A (15 mg/kg/day; Gel-Pharma) treatment starting the day following surgery, until euthanasia.

### Diffusion Tensor Imaging processing

DWI scans were pre-processed using FSL Diffusion Toolbox ^46^ of FSL 5.0.11 software. Each image was corrected for eddy currents distortions and head motion by affine registration to the average b0 image. A binary brain mask was obtained to remove non-brain tissue. The diffusion tensor model was adjusted to generate the FA and MD maps for all NHPs. Each FA and MD map was registered nonlinear to the macaque Rhesus template INIA19 ^47^. Region of interest (ROI) of bilateral anterior, medial and posterior putamen were drawn from the MRI template and the average of FA and MD for each NHP were computed to the corresponding ROI.

### PET acquisition and imaging analysis

To analyze the functional imaging outcome of cell transplantation, PET studies with a vesicular monoamine transporter probe were performed 9 months post-surgery. (+)-α-[^11^C]dihydrotetrabenazine (^11^C-DTBZ) was synthesized at the Unidad Radiofarmacia-Ciclotron of Universidad Nacional Autónoma de México, in the Tracerlab FXC-Pro synthesis module (GE Healthcare, Uppsala, Sweden) following published procedures ^48^. For each animal, initial anesthesia was induced with Tiletamine/Zolazepam (125/125 mg) at a dose of 2 mg/kg (i.m.). All animals were fasted overnight before the PET scan. Anesthesia was maintained during the scans with isoflurane (1-2 %, O_2_ flow rate of 2 l/min). PET imaging was performed on a Siemens Biograph PET-CT scanner at the Instituto Nacional de Neurología y Neurocirugía. All animals were fixed in the stereotaxic frame in a supine position with the head centered in the FOV and ^11^C-DTBZ was injected intravenously (74 ± 18.5 MBq). The parametric PET images were transformed into standard MRI space using a non-linear registration to analyze the ^11^C-DTBZ binding. The analysis was performed in bilateral striatal ROIs (left and right), which were manually segmented from the MRI template. Imaging analysis of the control and MPTP group (n = 3 for each) were averaged to obtain the mean binding for each group. For Sham, Grafted 1 and Grafted 2, ^11^C-DTBZ binding was obtained individually and values were transformed to z-score to normalize ^11^C-DTBZ binding.

### Microdialysis experiments and high-performance liquid chromatography (HPLC) analysis

Ten months after grafting, 28-mm long microdialysis probes manufactured by ourselves according to published methods ^49^. Membranes were introduced in both putamina to measure extracellular DA concentrations. The active part of the dialysis probe was a polyacrylonitrile membrane (molar weight cutoff, 40 kDa) and its inlet was connected to a syringe mounted on a microperfusion pump. *In vitro* recovery experiments with the dialysis membranes had values of 15-25% for DA and 3,4-dihydroxyphenylacetic acid (DOPAC). The probes were perfused with artificial cerebrospinal fluid at 2 μl per minute and fractions were collected every 12 minutes. Monoamines were stabilized by adding 0.1 N perchloric acid, 0.02% EDTA, and 1% ethanol. Extracellular DA increases were obtained through chemical depolarization (isosmotic solution containing 100 mM KCl) or 30 μM amphetamine and quantified for monoamine content by HPLC. No recovery correction was performed. After histological analysis, we found that all microdialysis probes were located in the putamina.

Twenty µL of dialysate samples were injected into the solvent stream of a HPLC system, using a reversed-phase column (C18, 3 μm; 2.1×50 mm) coupled to a pre-column (Atlantis, Waters®, U.S.) with a mobile phase solution containing sodium acetate 25 mM, EDTA 0.01 mM; citric acid 25 mM and 1-octanesulfonic acid 1 mM dissolved in milli-Q water and mixed with acetonitrile in a proportion of 95:5, respectively (pH of 3.35 ± 0.05 at a flow rate of 0.35 mL/min). DA and DOPAC detection were performed by a single-channel electrochemical detector (Waters® model 2465, U.S.) at 450 mV at a temperature of 30°C and quantified by peak height measurements against standard solutions. For cultures, samples were collected at day 60 of differentiation ^50^.

### Immunohistochemistry and postmortem cell count

NHPs were perfused via the right carotid with ice-cold saline followed by 4% paraformaldehyde (PFA). The brain was post-fixed overnight by immersion in 4% PFA and thereafter equilibrated in increasing sucrose solutions (10%, 20% and 30%) at 4°C. Tissue was sectioned on a cryostat in 20 μm slices that were serially recovered on individual slides. For immunohistochemical analyses, sections were permeabilized and blocked for 1 hour with 0.3% Triton X-100 and 10% normal goat serum in PBS. Samples were incubated overnight at 4°C with the primary antibodies diluted in PBS containing 10 % normal goat serum. Alexa-Fluor secondary antibodies were used diluted in PBS/10% normal goat serum for 1 hour. Nuclei were stained with Hoechst 33258. Immuno-reactive tissue was visualized using an epifluorescence microscope (Nikon eclipse TE2000-U) or a confocal laser microscope (Carl Zeiss LSM710). Negative controls without primary antibodies were included to confirm the specificity of detection. The number of TH+ in the *substantia nigra* and TH+/EGFP+ (in both putamen) neurons were quantified in Sham and Grafted NHPs. Twenty-four successive coronal sections throughout the grafts were photographed and the number of cells was estimated at 20X magnification.

### Statistical analysis

For cell cultures, behavioral assessment and microdialysis, we used unifactorial analysis of variance followed by Tukey’s *post hoc* analysis. The DTI results were plotted using the software GraphPad Prism program version 6. The Pearson’s correlation matrix between DTI measures and the behavioral outcome was computed in R 3.3.3 version. A value of *P* <0.05 was considered significant.

## Acknowledgments

We are grateful to M.Sc. César Augusto Rodríguez Balderas and D.V.M. Roberto Ramirez Hernández for taking care of the NHPs. Dr. Juan José Ortíz Retana of the Unit Magnetic Resonance of the Instituto de Neurobiología UNAM supported the acquisition of the structural and DTI scan data. We thank Dr. José Segovia (Cinvestav-Zacatenco) for providing *nu/nu* mice for teratoma formation. We acknowledge to Dr. Mabel Cerrillo and B.Sc. Nayelli Meza González of the laboratory “Reproducción y Genética AGN”, Hospital Ángeles del Pedregal in Mexico City for karyotype analysis. Histological stainings were made by the Histology Unit of Instituto de Fisiología Celular. Dr. Ruth Rincón helped with the acquisition of confocal images. We also thank Dr. M.A. Ávila-Rodríguez at the Unidad Radiofarmacia-Ciclotron, Facultad de Medicina, Universidad Nacional Autónoma de México in Mexico City, Sarahi Rosas and the Molecular Imaging PET-CT Unit of Instituto Nacional de Neurología y Neurocirugía for technical support in PET acquisition and imaging analysis. We thank Dr. Adriana Pacheco for facilitating sequencing equipment and assays on the EcoGenomics Laboratory at Tecnológico de Monterrey. This work was supported by PAPIIT, UNAM (IN213719 to I.V.), CONACYT-México: 222009 (A.C-R.), 256092 (I.V.), SALUD-272815 (I.V.) and 255747 (V.T.). V.R-M was supported by Instituto de Salud Carlos III-FEDER (CPII17/00032).

## Author contributions

A. L-O., I. E-A., J. F-R., A. C-R. and I. V. designed research. A. L-O., I. E-A., G. R-G., R. L-R., C. M-R., B. U-Ch., T. B-G., V. C-Ch., X. F-P., F. C., C. A. R., C. A., A. C-R. and I. V. performed experiments. N. K., L. R., L. R-V., V. T., J. B. and V. R-M. provided reagents, equipment and expertise. A. L-O., I. E-A., G. R-G., A. C-R. and I. V. drafted the manuscript. All authors approved the final version.

## Additional information

This manuscript contains Supplementary Figures and videos.

