## Supplemental file for "Human embryonic stem cells-derived dopaminergic neurons transplanted in parkinsonian monkeys recover dopamine levels and motor behavior"

**Supplementary Legends and Figures**

**Supplementary Figure 1.- Pluripotency of cell lines H9 and H9-EGFP. A)** Immunocytochemical analysis on H9 cell line. Scale bar, 200 µm. **B)** Gene expression, by RT-qPCR, for *SOX2*, *OCT4*, and *KLF4* on H9-EGFP cell line. Negative controls are BJ1 human fibroblasts. **C)** H9-EGFP cell line presents a normal female karyotype (46XX). **D)** Teratoma formation after H9-EGFP subcutaneous inoculation where cells representative of the three embryonic germ layers can be identified by hematoxylin and eosin staining (upper panel). The lower panel shows EGFP endogenous fluorescence. Mean ± SEM; $*P < 0.05$, n=3 independent experiments. Scale bar, 300 µm.

**Supplementary Figure 2.- Comparison of expression data at similar time points from previous work with our RNA-Seq results.** The microarray results from Kriks et al [12] were aligned with the reads of the same genes found in our sequencing experiments. Note that although the times are not exactly the same (day 13 vs day 14 and day 25 vs day 28), the patterns of expression are similar, indicating that our differentiation proceeded as reported for the floor-plate induction of dopamine neurons.

**Supplementary Figure 3. MPTP intoxication diminishes Fractional Anisotropy (FA) and increases Mean Diffusivity (MD) in the putamen when compared to healthy subjects. A**) MRI template and region of interest (ROI, in yellow) measured in each putamen onto anatomical brain image. FA and MD were determined bilaterally in the putamen. Three healthy NHPs were used for Control (non-parkinsonian) and 3 for the MPTP group. **B)** Graphs showing the comparison of FA and MD between the control and MPTP-treated groups, in the right and left putamina.

### Supplementary Figure 1

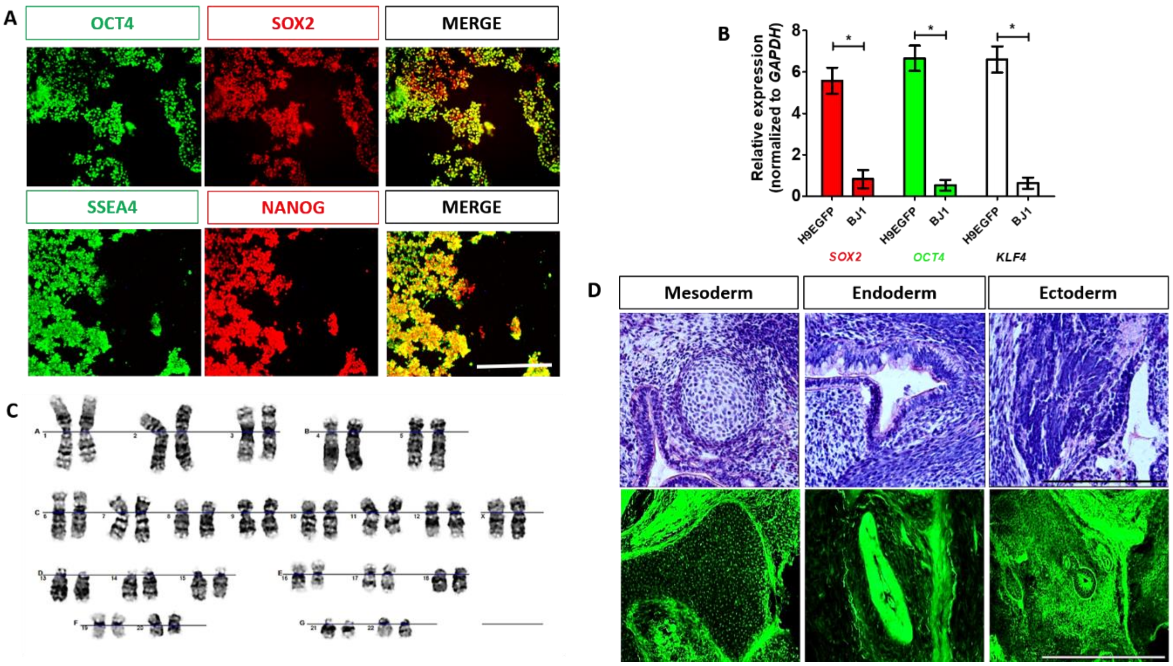

**Supplementary Figure 2**

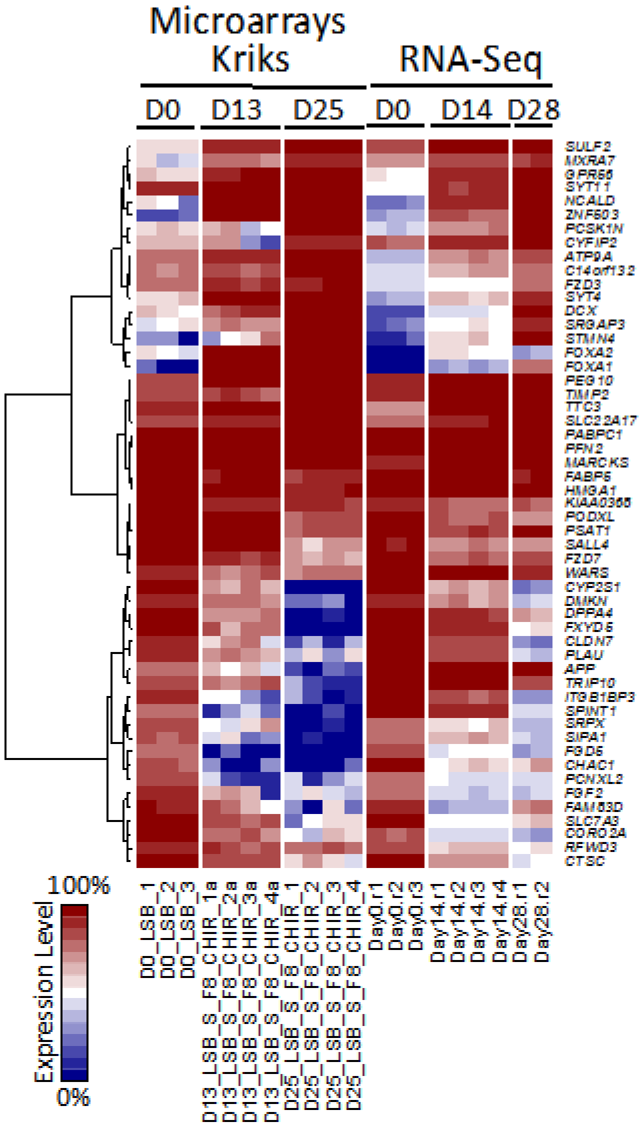

**Supplementary Figure 3**

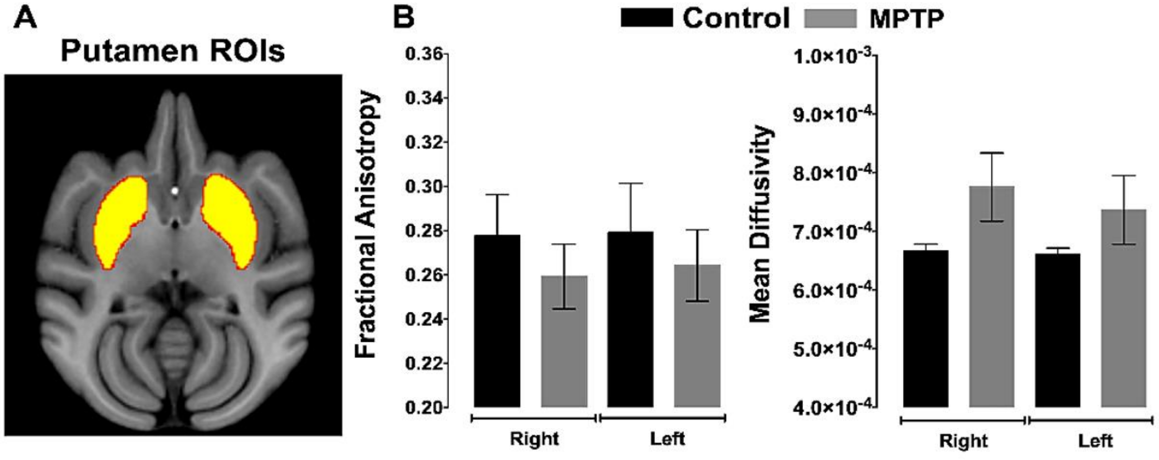
